# Probing Multiple Algorithms to Calculate Brain Age: Examining Reliability, Relations with Demographics, and Predictive Power

**DOI:** 10.1101/2022.06.17.496576

**Authors:** Eva Bacas, Isabella Kahhalé, Pradeep R Raamana, Julian B Pablo, Apurvaa S Anand, Jamie L Hanson

## Abstract

The calculation of so-called “brain age” has been an emerging biomarker in aging research. Data suggests that discrepancies between chronological age and the predicted age of the brain may be predictive of mortality and morbidity (for review, see Cole, Marioni, Harris, & Deary, 2019). However, with these promising results come technical complexities of how to calculate brain age. Various groups have deployed methods leveraging different statistical approaches, often crafting novel algorithms for assessing this biomarker. There remain many open questions about the reliability, collinearity, and predictive power of different algorithms. Here, we complete a rigorous systematic comparison of three commonly used, previously published brain age algorithms (XGBoost, brainageR, and DeepBrainNet) to serve as a foundation for future applied research. First, using multiple datasets with repeated MRI scans, we calculated two metrics of reliability (intraclass correlations and Bland–Altman bias). We then considered correlations between brain age variables, chronological age, biological sex, and image quality. We also calculated the magnitude of collinearity between approaches. Finally, we used canonical regression and machine learning approaches to identify significant predictors across brain age algorithms related to clinical diagnoses of mild cognitive impairment or Alzheimer’s Disease. Using a large sample (*N=2557*), we find all three commonly used brain age algorithms demonstrate excellent reliability (r>.9). We also note that brainageR and DeepBrainNet are reasonably correlated with one another, and that the XGBoost brain age is strongly related to image quality. Finally, and notably, we find that XGBoost brain age calculations were more sensitive to the detection of clinical diagnoses of mild cognitive impairment or Alzheimer’s Disease. We close this work with recommendations for future research studies focused on brain age.

## INTRODUCTION

Aging is a process that involves multiple factors, with the causes of aging still poorly understood. As the human lifespan has increased, an increased proportion of the global population is beginning to show age-associated functional declines and disease (Roser et al., 2013). It is therefore critical for us as a society to delineate the pathways more richly to longevity, as well as age-associated deteriorations. To understand and predict these age-related differences in health and disease, novel approaches have emerged attempting to quantify an individual’s “biological age” and its divergence from chronological age. These have included examination of telomeres, epigenetics, and other molecular and cellular elements (Aubert & Lansdorp, 2008; Fraga & Esteller, 2007; Mather et al., 2011; Pal & Tyler, 2016). Moving beyond molecular and cellular assays, calculation of so-called “brain age” is a novel biomarker that has recently emerged in aging research. This type of biological age is estimated from neuroimaging scans using data science and machine learning algorithms. Given the brain’s central role in regulating behavior and multiple neuroendocrine processes, metrics of brain age may be a particularly predictive indicator of age-related mortality and morbidity.

Surveying work on brain age to date, different research groups have focused on the discrepancies between brain age and chronological age. In this work, we refer to the difference between predicted brain age and chronological age as *brain age delta*, adopting the terminology used in Bashyam et al. (2020). Of note, other scholars use varying terms to refer to this difference, including brain age gap and brain-PAD (Kaufmann et al., 2019; Cole et al., 2018). This approach presumes that larger differences reflect poorer health, and suggestive evidence is growing to validate this idea. For example, Cole and colleagues (2018) found that greater brain age delta was associated with a number of physiological measures related to senility, including weaker grip strength, poorer lung function, and longer times to walk a short distance. Particularly notable, greater brain age delta – derived from MRI scans of participants in their early 70s – was related to mortality years later. In this sample, older brain age was associated with reduced lifespan, with each additional year of brain aging being related to a ∼6% increase in the likelihood of death between the ages of 72 and 80. In addition to aspects of senility and mortality, brain age has also been connected to several psychiatric and neurodegenerative conditions. Patients with serious psychiatric and neurological disorders show increased brain age, including patients with schizophrenia (Shahab et al., 2019), depression (Kaufmann et al., 2019), borderline personality disorder (Koutsouleris et al., 2014), and Alzheimer’s disease (Franke & Gaser, 2012, 2019; Gaser et al., 2013; Ly et al., 2020).

While promising, research deploying these approaches is still in its infancy. There are multiple unique algorithms to calculate brain age developed by pioneering groups. In previous brain age research, it is common to develop and apply newly developed algorithms in the same research report. From a technical perspective, the calculation of brain age uses a wide variety of machine learning approaches, including Gaussian process regression, regularizing gradient boosting, and more recently, deep learning models. This has led to a host of debates in the field about how to conceptualize and validate these different algorithms (see Bashyam et al., 2021; Hahn et al., 2021). Adding to this complexity, the diverse aims and applications of brain age studies result in a variety of benchmarks. For example, one could examine the correlation between brain age and chronological age, different estimates of error in prediction (i.e., mean squared error), or the ability of algorithms to identify different age-related conditions and declines (e.g., early signs of Alzheimer’s disease). Examined collectively, many open questions exist regarding the strengths and limitations of different algorithmic derivations of brain age.

Here, our aim was to provide a systematic comparison of three major brain age algorithms to serve as a point of reference for future applied research. To compare the algorithms, we looked at predictive power, reliability, and noise sensitivity. As previous research has examined relationships between brain age delta and age-related decline, we focused on predictive power as it relates to diagnoses of mild cognitive impairment (MCI) and Alzheimer’s disease (AD). Reliability is a relevant point of comparison for researchers working with longitudinal data, and noise sensitivity is important for researchers working with noisy data or populations that typically produce noisy data (e.g. neuropsychiatric patients, children) (Wylie et al., 2014).

Related to intra-algorithm reliability, we use intraclass correlation and Bland–Altman bias metrics to compare algorithmic performance across repeated MRI scans of the same individuals. We also examined correlations between different algorithms to see the reliability of results across approaches. To understand the influence of noise in MRI images, we examined relations between image quality and brain age calculation. We examined reliability and noise sensitivity in an older as well as a young sample of participants. This was motivated by the fact that brain age algorithms are typically developed in older age samples, but now are being applied to young participants (e.g. Keding et al., 2021). Finally, related to predictive power, we used a data-driven, machine learning approach to identify significant predictors across brain age algorithms related to a commonly used outcome, clinical diagnoses of MCI or Alzheimer’s Disease.

## METHODS

### Datasets

For our analyses, we examined three large open-access MRI datasets that included participants 19-100 years of age (*Analytic N=2557*). All data were collected on 3-Tesla MRI scanners. Since our project aimed to examine the reliability of brain age metrics, we leveraged two datasets with repeated MRI scans (Amsterdam Open MRI Collection [AOMIC]; Open Access Series of Imaging Studies [OASIS]). We also examined a third dataset that had reasonable variability in age, but only one MRI scan (Human Connectome Project-Aging [HCP-A]). A major concern in the development of statistical models is the issue of overfitting, when a model performs well on the training data but fails to generalize to new data (Ying, 2019). To account for potential overfitting, we specifically selected datasets that were not used in training data for any of the three brain age algorithms compared in our analysis.

Basic descriptions of the datasets are found in the following section. Additional details of the data collection are summarized in our supplemental materials.

#### Amsterdam Open MRI Collection (AOMIC)

AOMIC is an open-access neuroimaging dataset including structural and functional MRI scans (Snoek et al., 2021). Here, we analyze the “ID1000” subset of the data which comprised healthy young adults aged 19-26 (*N = 928*) scanned between 2010 and 2012 at University of Amsterdam. Participants were recruited from the general Dutch population, with efforts to recruit from a variety of educational backgrounds. Each participant was scanned three times in a single session using the same imaging parameters. Specifically, MR images were acquired with a Phillips Intera 3T scanner. T1-weighted MR images were acquired using a sagittal 3D-MPRAGE sequence. In addition to MRI scanning, participants also completed many well-validated self-report scales and behavioral assessments, including measures of cognitive ability, personality, and motivation. Additional details of the data collection are summarized in our Supplemental Materials (Table S1).

#### Open Access Series of Imaging Studies (OASIS)

OASIS is a multimodal neuroimaging project centralized at Washington University in St. Louis (LaMontagne et al., 2019). For our work, we used OASIS-3, a dataset of normally aging and Alzheimer’s disease patients (starting N = 1098) aged 42-95. Six-hundred and five of these participants were neurologically healthy, while 493 participants presented with mild-cognitive impairment, Alzheimer’s disease, and other neurological conditions of concern. Participants were recruited from other ongoing projects at Washington University in St. Louis focused on Alzheimer’s and aging. For our work, we used data collected on Siemens TIM Trio 3T scanners. A subset of OASIS were scanned at multiple timepoints months or years apart (“sessions”) and some participants were scanned multiple times in a single session (“runs”). We included only participants with multiple runs per session. This eliminated 329 participants.

#### Human Connectome Project in Aging (HCP-A)

HCP-A is an ongoing project collecting MRI data from 1200 healthy adults greater than 36 years of age (Bookheimer et al., 2019). Participants were recruited using flyers, advertisements, and outreach at community centers, with efforts to recruit a balanced number of participants across socioeconomic classes. Data collection was completed at 4 research sites: Massachusetts General Hospital, University of California at Los Angeles, University of Minnesota, Washington University in St. Louis. For this work, our sample was composed of 725 participants aged 36-100. MR images at all sites were collected using matched Siemens Prisma 3T scanners. T1-weighted MR images were collected using a multi-echo sagittal 3D-MPRAGE sequence. Participants also completed many well-validated self-report scales and behavioral assessments, including measures of cognitive ability, personality, and mental health.

### Assessment of MRI Image Quality

Past work from our research group (Gilmore et al., 2021) has found that T1-weighted image quality is related to volumetric measures from commonly used morphometric tools suites (e.g. Freesurfer). We therefore assessed image quality to: 1) exclude particularly high-motion scans; and 2) investigate the impact of image quality on brain age estimates. To assess MRI quality, we generated a quantitative metric (“CAT12 score”) using the Computational Anatomy Toolbox 12 (CAT12). This metric considers four summary measures of image quality: noise-to-contrast ratio, coefficient of joint variation, inhomogeneity-to-contrast ratio, and root-mean-squared voxel resolution. CAT12 normalizes and combines these measures using a kappa statistic-based framework. The score is a value from 0 to 1, with 0 being the lowest quality and 1 being the highest quality. Informed by our past work (e.g., Gilmore et al., 2021), we excluded all lower quality scans, specifically with CAT12 scores <0.8.

### Analytic Sample and Brain Age Algorithms

After removing data with quality and preprocessing issues, our total analytic sample was 2557 participants, with 928 subjects from AOMIC, 705 subjects from HCP-A, and 584 subjects from OASIS.

Due to their use in recent brain age publications, we tested three brain age algorithms on our dataset: Kaufmann et al. (2019) (referred to as “XGBoost”), Cole et al. (2018) (referred to as “brainageR”), and Bashyam et al. (2020) (referred to as “DeepBrainNet”). We selected these three algorithms based on popularity amongst researchers and open access code. DeepBrainNet and brainageR operate on raw T1-weighted MRI scans, and XGBoost requires preprocessing using Freesurfer (Fischl, 2012), an open source MRI processing software package. We provide brief summaries of the algorithms below. For detailed descriptions of model structure, please see the original papers cited here. Below, we briefly outline the varying benchmarks used in original publications, noting correlations between brain age and chronological age when available. Of note, past work has used many different statistical tests and validation approaches (e.g. mean squared error, accuracy), making exact comparisons between algorithms inaccurate or not possible.

#### XGBoost (Kaufmann et al., 2019) Brain Age Algorithm

XGBoost uses gradient tree boosting to predict brain age based on 1118 features extracted using Freesurfer. These features consist of thickness, area, and volume measurements from a multimodal parcellation of the cerebral cortex, cerebellum, and subcortex. Relevant code is available at: https://github.com/tobias-kaufmann/brainage. This algorithm was trained on a large and diverse sample (N = 39,827, female = 18,990). The sample was made up of healthy controls aged 3-89 drawn from 42 different datasets. To account for potential variation, Kaufmann et al. trained separate models for male and female brain age. We deployed this algorithm by first completing standard processing approaches in Freesurfer 7.1 (http://surfer.nmr.mgh.harvard.edu). The technical details of this software suite are described in prior publications (Dale et al., 1999; Fischl et al., 1999, 2002, 2004). Briefly, this processing includes motion correction and intensity normalization of T1-weighted images, removal of non-brain tissue (Ségonne et al., 2004), automated Talairach transformation, segmentation of white matter and gray matter volumetric structures, and derivation of cortical thickness. Freesurfer processing was implemented via Brainlife.io (brainlife/app-freesurfer), which is a a free, publicly funded, cloud-computing platform for developing reproducible neuroimaging processing pipelines and sharing data (Avesani et al., 2019; Pestilli, 2018).

#### brainageR (Cole et al., 2018) Brain Age Algorithm

brainageR uses Gaussian Process Regression to predict brain age based on raw, unprocessed, T1-weighted MR images. Relevant code is available at: https://github.com/james-cole/brainageR. This software uses SPM12 for segmentation and normalization with custom brain templates, and loads these images into R using the RNfiti package. Gray matter, white matter and CSF vectors are then used to predict a brain age value with a model previously trained with kernlab. This algorithm was trained on a sample (*N = 2001*) of healthy adults aged 18-90, including scans from 14 different studies. After model building and tuning, Cole et al. found brain age and chronological age were correlated for r=0.92.

#### DeepBrainNet (Bashyam et al., 2020) Brain Age Algorithm

DeepBrainNet is a 2D Convolutional Neural Network (CNN) built using the inception-resnetv2 framework. Notably, this model was initialized with random weights and trained exclusively on MRIs to create a brain-specific model. With this algorithm, raw, unprocessed, T1-weighted MR images are n4 bias corrected, skull-stripped, and affine registered to an MNI-template. This algorithm was implemented through the ANTsRNet package, an implementation of Advanced Normalization Tools (ANTs) in the R programming language (Tustison et al., 2021). Relevant code for this algorithm is located here: https://github.com/ANTsX/brainAgeR. This algorithm was originally trained on a sample on a sample (N = 11729) of healthy controls aged 3-95 drawn from 18 different datasets. Bashyam and colleagues found a correlation of r=0.978 between predicted brain age and chronological age; however, those authors purposefully selected a “*moderately fit*” model over a loosely or tightly fit model. This was motivated by the belief that a moderately fit model would better reveal individual differences in pathology.

### Statistical Analyses Related to Brain Age Reliability

To assess the reliability of brain age calculation by algorithm, we used two approaches of looking at reliability: intraclass correlation coefficient (ICC) and Bland-Altman analysis. Of note, for these analyses, we only used data from AOMIC and OASIS due to the repeated scans for each participant. ICC is a descriptive statistic indicating the degree of agreement between two or more sets of measurements. The statistic is similar to a bivariate correlation coefficient insofar as it has a range from 0-1 and higher values represent a stronger relationship. An ICC differs from the bivariate correlation in that it works on groups of measurements and gives an indication of the numerical cohesion across the given groups (McGraw & Wong, 1996). We calculated ICCs using the statistical programming language R, with the icc function from the package *‘irr’* (Gamer et al., 2012). A two-way model with absolute agreement was used in order to investigate the exact estimate of brain age for each repeated scan. Although there are no definitive guidelines for precise interpretation of ICCs, results have frequently been binned into three (or four) quality groups where 0.0-0.5 is “*poor*”, 0.50-0.75 is “*moderate*”, 0.75-0.9 is “*good*” and 0.9-1.0 is “*excellent*” (Cicchetti, 1994; Koo & Li, 2016). Additionally, Bland-Altman analyses investigate reliability by considering the differences between paired groups of measurements. In our analysis, the paired groups are brain age predictions across two scans. As AOMIC contains three scans per participant, we compared across all pairings of scans (e.g., scan 1 versus scan 2, scan 2 versus scan 3, and scan 1 versus scan 3) and took the average difference metric. In addition to these raw difference scores (i.e., the difference across two instances of measurement), we also considered the proportion of difference, calculated by taking the difference divided by the mean value for a given pairing of measurements.

### Statistical Analyses Connected to Image Quality and Sociodemographic Factors

After reliability analyses, we reduced our analytic sample to include only the highest quality scan from each participant. This was done to maintain statistical independence of observations. Put another way, including repeated scans in standard regression models, but presuming independence, would violate fundamental statistical assumptions. The highest quality MRI scan was selected using CAT12 scores. Using these higher quality scans, we calculated bivariate correlations between brain age and real age. We also examined relations between brain age and multiple relevant variables, including image quality, and participant sex. In these analyses, we looked at associations with both brain age and brain age delta in all three of our datasets. Of note, many brain age researchers choose to correct for age-related bias using linear models or other methods (e.g., (Le et al., 2018). This correction is often done in relation to group or individual difference variables. However, given that we were not focused on these differences and instead wanted to understand the full effects of potential confounds on the algorithms tested, we used uncorrected correlations in this portion of our analyses.

### Statistical Prediction Using Brain Age Variables

We used the OASIS dataset to look at the association between brain age variables and MCI (MCI) and/or Alzheimer’s disease. Based on previous findings, we would expect a greater brain age delta to be related to neurodegenerative conditions such as Alzheimer’s disease. To investigate this hypothesis, participants were coded for presence of MCI and/or Alzheimer’s Disease (presence=1; absence=0). Individual logistic regression models were fit for each algorithm. Presence of MCI and/or Alzheimer’s Disease status were entered as the dependent variable. Brain age delta, chronological age, sex, and CAT12 scores were included as independent variables. Finally, given potential collinearity between brain age algorithms, we also fit an elastic net (EN) model to determine the best combination of predictors of clinical status. In brief, EN machine learning algorithms use both different regression penalties (i.e., lasso [L1]; ridge [L2]) to prevent overfitting of the model, compromising between penalties by weighting the proportion of ridge and lasso penalties (α). To tune the penalty, cross-validation identifies a second parameter, λ, which is the magnitude of the shrinkage penalty. This modeling and parameter identification was implemented using the *beset* package in R (Shumake, 2021). This R library tests over 100 possible values of λ (autogenerated by the package), as well as three values of α: 0.01 (weighting the ridge penalty more heavily and therefore including more variables, 0.99 (weighting the lasso penalty more heavily and including fewer variables), and 0.50 (equal weighting of both penalties). Similar to above, the presence of Mild-cognitive impairment and/or Alzheimer’s Disease status was coded as a binary variable and entered as the dependent variable (Yes to either Mild-cognitive impairment and/or Alzheimer’s Disease = 1; No 0). Potential independent variables included brain age deltas from all three algorithms, chronological age, sex, and CAT12 scores. Nested ten-fold cross validation was performed and we output a summary ranking relative importance of the predictor variables.

## RESULTS

### Brain Age Reliability by Algorithm

Given that brain age algorithms are being used in older (Cole et al., 2018), as well as younger (Keding et al., 2021) samples, we computed test-retest reliability in two different open-access neuroimaging projects with repeated scans. For the young adults in the AOMIC project, all three algorithms obtained ICCs greater than 0.9 (XGBoost: r = 0.935, brainageR: r = 0.983, DeepBrainNet: r = 0.979). The mean proportion of difference across all three scans was small. XGBoost had a mean of 6.535%, followed by brainageR with 2.449% and DeepBrainNet with 2.04%. These differences are depicted in Figure 2. The mean difference across all three scans was similarly small. XGBoost and brainageR had slightly negative mean differences of -0.139±2.376 years and -0.098±0.777 years, and DeepBrainNet had a mean difference of 0.011±0.791 years.

**Figure 1.**
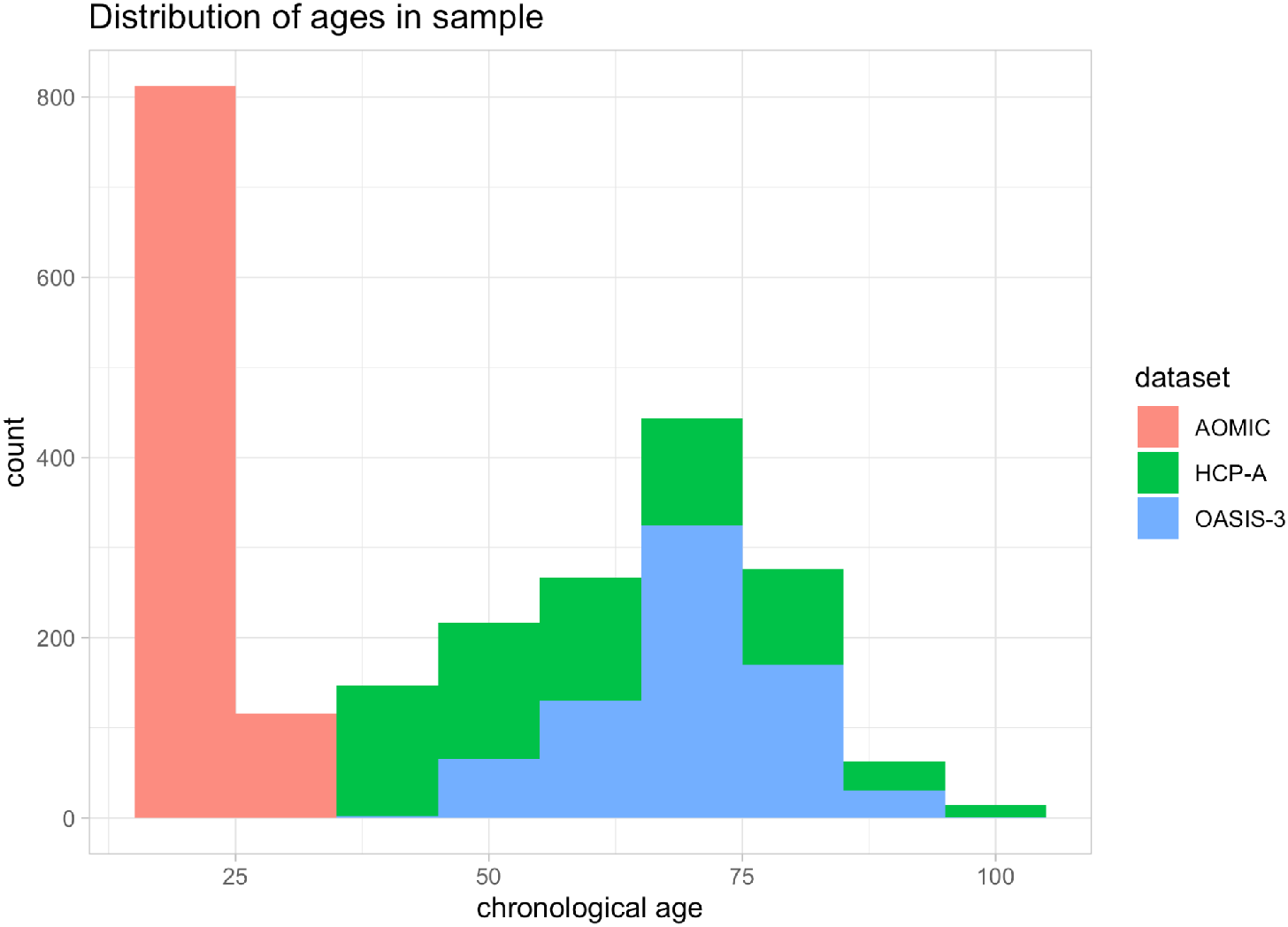
Distribution of participant age for the different projects we leveraged in our analyses. The horizontal axis depicts participant age in years, while the vertical axis shows the number of participants within a given age bin. Each dataset is shown in a different color, with AOMIC shown in red, HCP-A shown in green, and OASIS-3 shown in blue.

**Figure 2.**
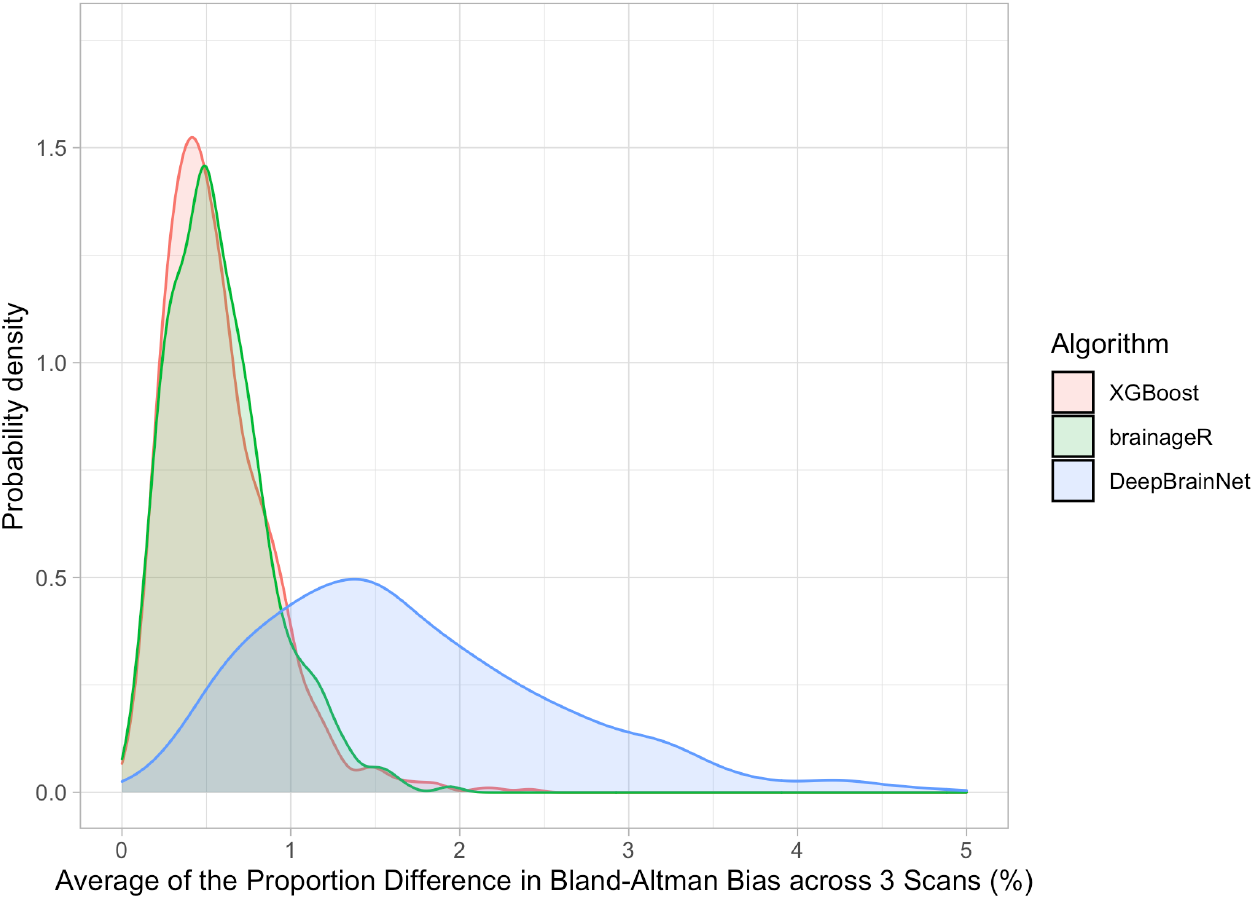
Density Plot of Differences in 3 Brain Age Algorithms across the AOMIC sample. The horizontal axis shows the portion of differences between repeated scans (as a percentage). The vertical axis is the density (or frequency) of such bias. Each brain age algorithm is shown in a different color with XGBoost shown in light red, brainageR shown in light green, and DeepBrainNet shown in light blue.

For the older adult sample in OASIS, all three algorithms obtained ICCs greater than 0.95 (XGBoost: r = 0.972, brainageR: r = 0.99, DeepBrainNet: r = 0.992). Mean proportion of difference and mean difference across both scans were again small, with XGBoost showing a mean proportion of difference of 2.397%, followed by brainageR with 1.518% and DeepBrainNet with 1.259%. These differences are depicted in Figure 3. XGBoost had a small positive mean difference of 0.021±1.865 years, and brainageR and DeepBrainNet had negative mean differences of -0.158±1.483 and -0.086±1.135. Examined collectively, all three algorithms obtained high ICCs and low mean differences for both AOMIC and OASIS data, suggesting that these algorithms are highly reliable.

**Figure 3.**
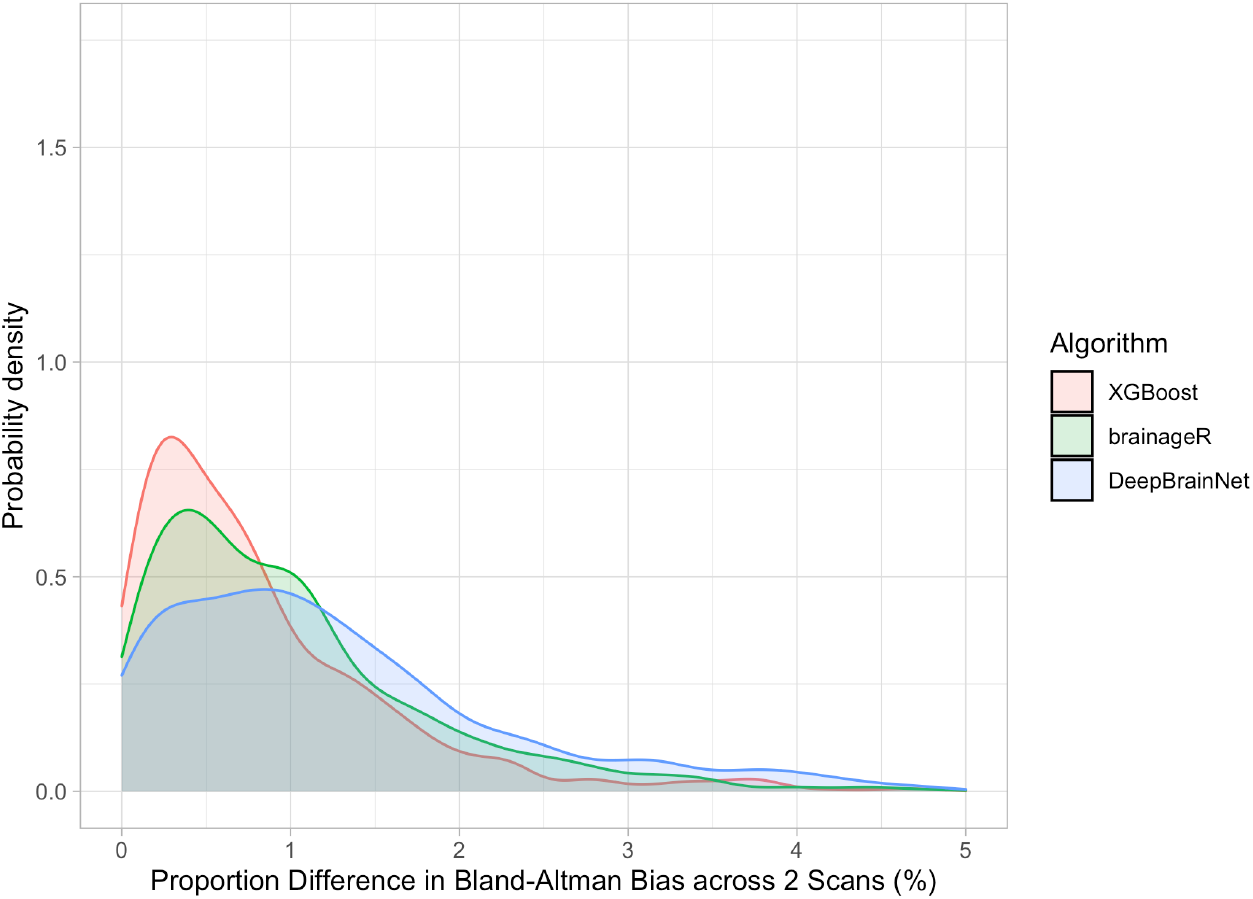
Density Plot of Differences in 3 Brain Age Algorithms across the OASIS sample. The horizontal axis shows the portion of differences between repeated scans (as a percentage). The vertical axis is the density (or frequency) of such bias. Each brain age algorithm is shown in a different color with XGBoost shown in light red, brainageR shown in light green, and DeepBrainNet shown in light blue.

Figures 2 and 3 show density plots of the mean proportion of differences for both AOMIC and OASIS. The x-axes represent the proportion difference, ranging from 0 to 5, and the y-axes represent the probability density, calculated using a Gaussian kernel.

### Relations Between Brain Age, Chronological Age, and Sex Brain Age Algorithms

When examining brain age and chronological age, there were strong correlations between these variables for each algorithm. All correlations were greater than 0.9, with r = 0.936 for XGBoost, r = 0.966 for brainageR, and r = 0.96 for DeepBrainNet. As Kaufmann et al. separated models for male and female subjects, we also computed correlations for brain age and chronological age for each sex. With XGBoost, correlations were comparable across sex (female: r = 0.932; male: r = .941). This pattern was similar for brainageR (female: r = 0.966; male: r = 0.968) and DeepBrainNet (female: r = 0.958; male: r = 0.963). All correlations had p-values < 0.001.

We additionally investigated associations between brain age delta and chronological age. Across the algorithms, there was a great variability in the relationship between brain age delta and chronological age: r = -0.776 for XGBoost, r = -0.225 for brainageR, and r = -0.489 for DeepBrainNet. All these correlations had p-values < 0.001.

All algorithms had significant negative correlations between brain age and image quality: XGBoost (r = -0.381, p < 0.001), brainageR (r = -0.458, p < 0.001), DeepBrainNet (r = -0.464, p < 0.001).

In comparison to brain age and image quality, the relationship between brain age delta and image quality showed great variability in correlations. For XGBoost, there was modest correlation between image quality and brain age delta (r = 0.36, p < 0.001). brainageR also had a small negative correlation between brain age delta and quality (r = -0.084, p < 0.001). For DeepBrainNet, there was no significant relationship between brain age delta and image quality (r = 0.033, p = 0.105).

### Correlation Variability by Dataset

Of important note, the magnitude of correlations between brain age and chronological age varied significantly across datasets. In the younger AOMIC sample, brainageR performed the best with a modest correlation of 0.44 (p < 0.001), followed by DeepBrainNet (r = 0.365, p < 0.001) and XGBoost (r = 0.303, p < 0.001). The correlation between brain age delta and real age was less than 0.1 for all algorithms; of note, this dataset has a very narrow age range, with the majority of participants in their 20s. The correlations between image quality and brain age delta was also less than 0.1 for all algorithms.

Interestingly, in the older samples (OASIS and HCP-A), there was a significant jump in the magnitude of the correlation between brain age and chronological age. For HCP-A, all three algorithms show correlations greater than 0.8 for brain age and real age, with brainageR again the highest (r = 0.917, p < 0.001), followed by DeepBrainNet (r = 0.903, p < 0.001) and XGBoost (r = 0.835, p < 0.001). HCP-A features the largest age range of all three datasets used in our analyses. The effect of age on brain age delta was significant for all algorithms: XGBoost had a relatively strong negative correlation (r = -0.768, p < 0.001), followed by brainageR (r = -0.213, p < 0.001) and DeepBrainNet (r = -0.169, p < 0.001). XGBoost additionally has a significant correlation of 0.359 (p < 0.001) between image quality and brain age delta.

DeepBrainNet performed the best on OASIS (r = 0.814, p < 0.001), followed by brainageR (r = 0.799, p < 0.001) and XGBoost (r = 0.733, p < 0.001). Similar to HCP-A, all three algorithms had significant correlations between age and brain age delta. XGBoost had the strongest correlation (r = -0.576, p < 0.001), followed by DeepBrainNet (r = -0.36, p < 0.001) and brainageR (r = -0.172, p < 0.001).

### Brain Age Delta as a Predictor of Dementia Status

In line with past work, individual logistic models found that brain age delta was a significant predictor of dementia status for XGBoost, brainageR, and DeepBrainNet. Chronological age and CAT12 score were additional significant predictors for all three models. Sex was not significant for any model. For XGBoost, brain age delta was a significant predictor (β = 0.207, z = 7.126, p < 0.001), along with real age (β = 0.134, z = 6.007, p < 0.001), and CAT12 score (β = -15.323, z = -3.495, p < 0.001). We found similar results for brainageR (brain age delta: β = 0.106, z = 5.064, p < 0.001; real age: β = 0.050, z = 3.244, p < 0.001; CAT12 score: β = -15.050, z = -3.496, p < 0.001) and DeepBrainNet (brain age delta: β = 0.178, z = 5.945, p < 0.001; real age: β = 0.097, z = 4.785, p < 0.001; CAT12 score: β = -13.110, z = -3.062, p < 0.001).

The results are summarized in figure 6 and tables 1, 2, and 3.

**Table 1.** A table depicting statistical parameter estimates from our logistic model (with clinical status as a binary dependent variable, and XGBoost brain age delta and other covariates as independent variables). Mean Parameter estimate, standard error, z-score, and Frequentist p-value are shown in different columns, with statistics for each independent variable shown in each row.

|  | XGBoost |  |  |  |
| --- | --- | --- | --- | --- |
|  | Estimate | Std. Error | z value | Pr(>\|z\|) |
| **Brain age delta** | 0.207 | 0.029 | 7.126 | <0.001 |
| **Real age** | 0.134 | 0.022 | 6.007 | <0.001 |
| **Sex - male** | 0.123 | 0.251 | 0.491 | 0.623 |
| **CAT12 score** | -15.323 | 4.385 | -3.495 | <0.001 |

**Table 2.** A table depicting statistical parameter estimates from our logistic model (with clinical status as a binary dependent variable, and brainageR brain age delta and other covariates as independent variables). Mean Parameter estimate, standard error, z-score, and Frequentist p-value are shown in different columns, with statistics for each independent variable shown in each row.

| brainageR |  |  |  |  |
| --- | --- | --- | --- | --- |
|  | Estimate | Std. Error | z value | Pr(>\|z\|) |
| **Brain age delta** | 0.106 | 0.021 | 5.064 | <0.001 |
| **Real age** | 0.050 | 0.015 | 3.244 | 0.001 |
| **Sex - male** | 0.216 | 0.247 | 0.876 | 0.381 |
| **CAT12 score** | -15.050 | 4.305 | -3.496 | <0.001 |

**Table 3.** A table depicting statistical parameter estimates from our logistic model (with clinical status as a binary dependent variable, and DeepBrainNet brain age delta and other covariates as independent variables). Mean Parameter estimate, standard error, z-score, and Frequentist p-value are shown in different columns, with statistics for each independent variable shown in each row.

| DeepBrainNet |  |  |  |  |
| --- | --- | --- | --- | --- |
|  | Estimate | Std. Error | z value | Pr(>\|z\|) |
| **Brain age delta** | 0.178 | 0.030 | 5.945 | <0.001 |
| **Real age** | 0.097 | 0.020 | 4.785 | <0.001 |
| **Sex - male** | 0.252 | 0.245 | 1.028 | 0.304 |
| **CAT12 score** | -13.110 | 4.281 | -3.062 | 0.002 |

**Figure 4.**
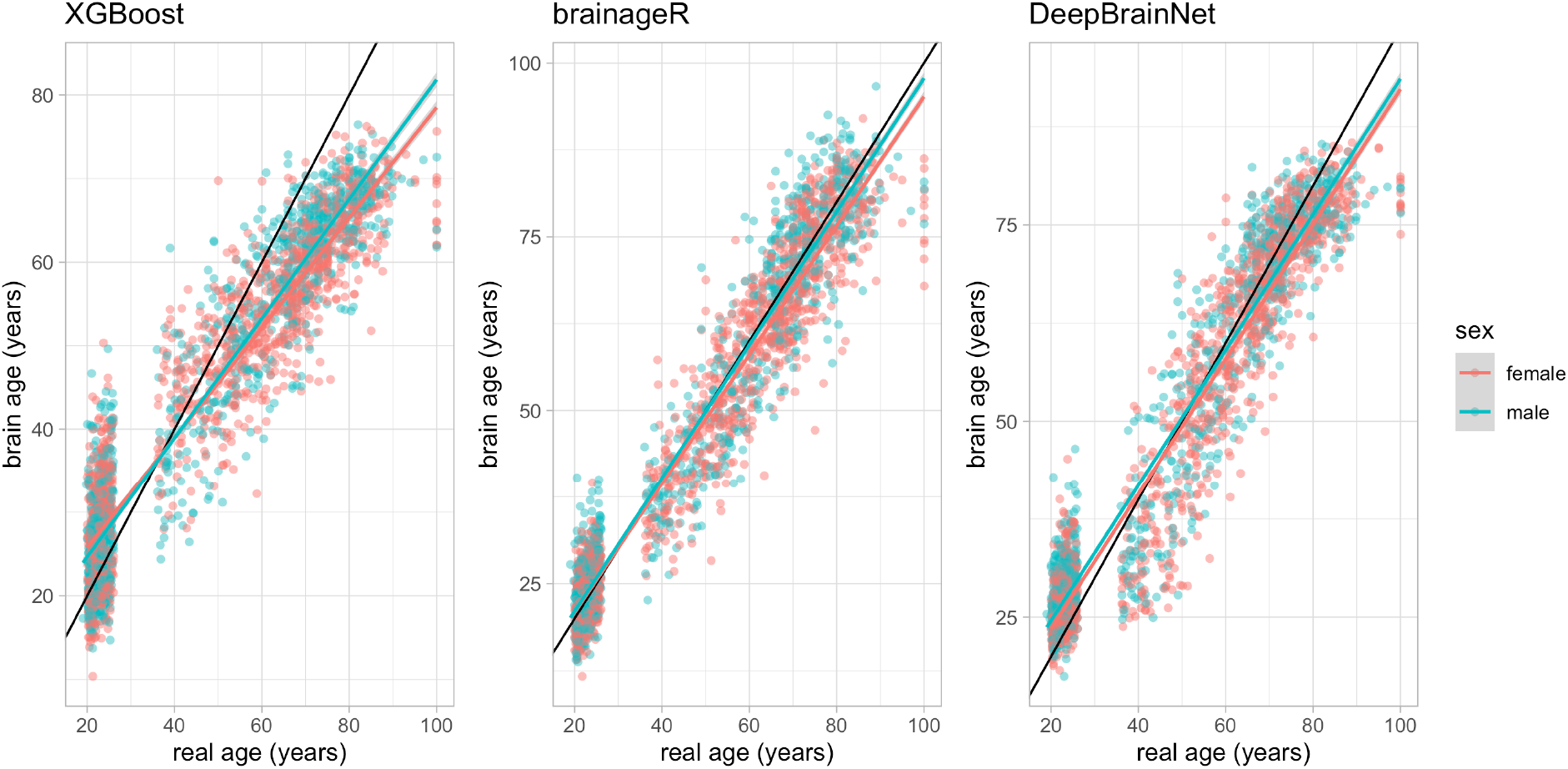
Scatterplots of the relationship between brain age and real age across all algorithms. There are three panels, each representing a different brain age algorithm—XGBoost is on the far-left panel, brainageR is in the middle, and DeepBrainNet is on the far-right panel. The horizontal axis shows participant chronological (real) age, while the vertical axis represent predicted brain age. In each panel, red dots represent female participants, and teal dots represent male participants. Of note, there are variation in the y-axes due to variation in predictions.

**Figure 5.**
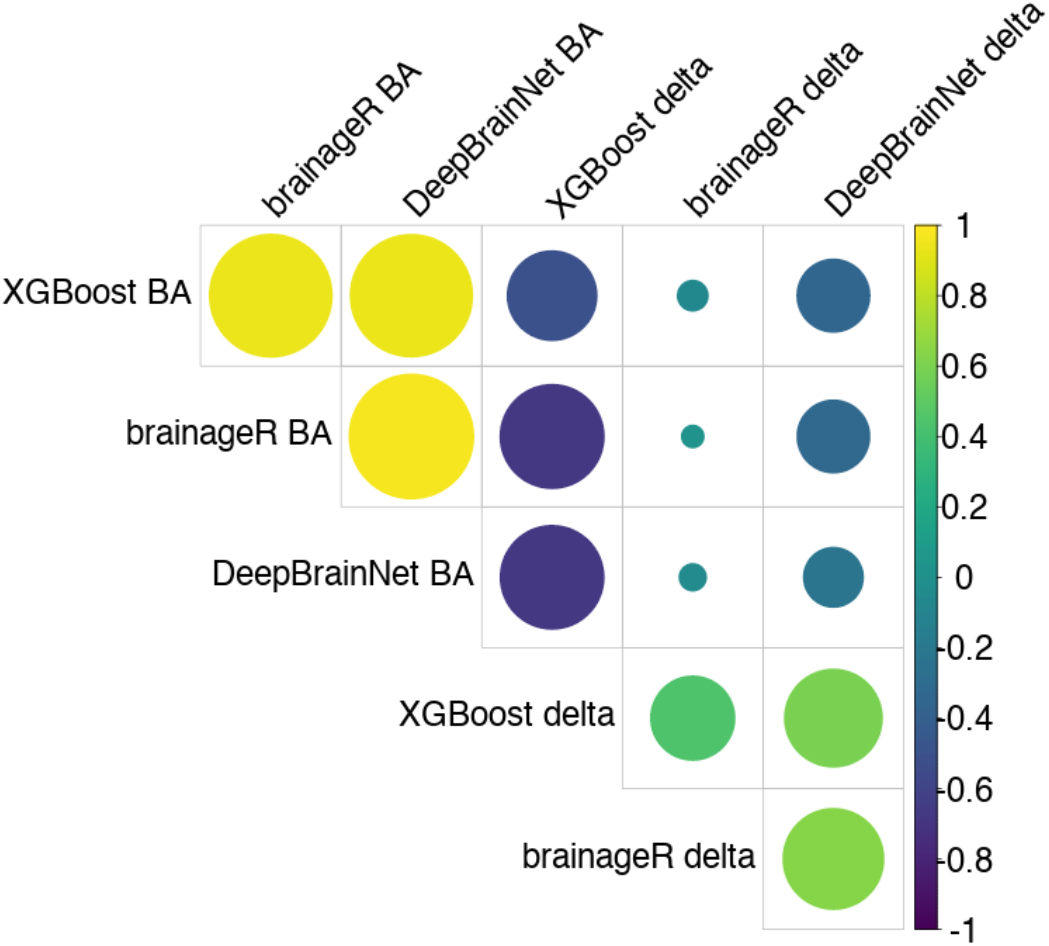
Correlation plot between brain age and brain age delta across algorithms. There are rows and columns representing different relevant variables. The correlation between variables is shown at the confluence of a row and a column. The strength of a correlation is represented by both the color and size of the dots. For example, larger yellow circles are high positive correlations, while larger blue circles are high negative correlations. Positive correlations are shown in yellow and green, while negative correlations are blue-green and darker blue. Of note, the abbreviation BA is for brain age.

**Figure 6.**
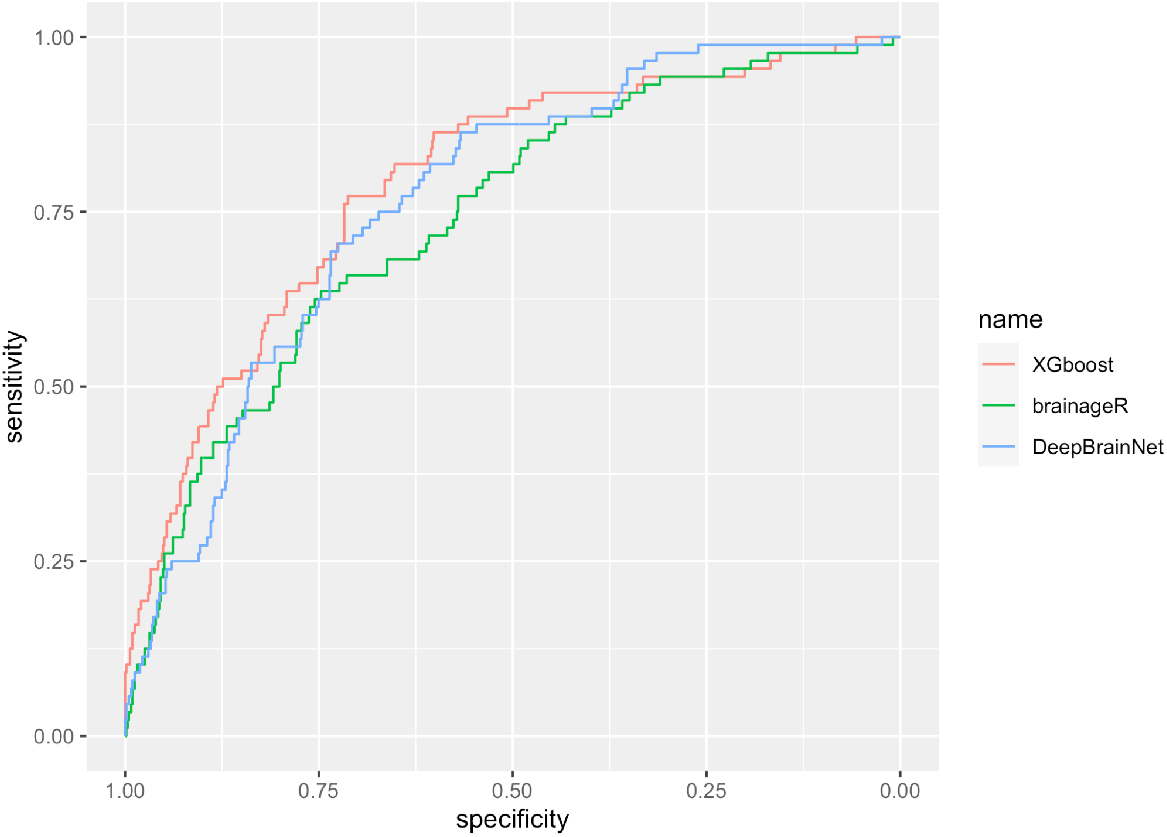
Receiver-operating characteristic curve for logistic models. These analyses were completed in the OASIS dataset where information about clinical status (Mild Cognitive Impairments or Alzheimer’s’ status) was available. Each algorithm is represented by a different color line, with XGBoost shown in light red, brainageR shown in light green, and DeepBrainNet shown in light blue. The horizontal axis shows the specificity (true negative rate), while the vertical axis shows the sensitivity (true positive rate) of each algorithm in detecting the presence of disease.

For the EN model, all variables were significant. The variables ranked by relative importance can be found in table 4. Chronological age is the most important coefficient. Of the brain age deltas, XGBoost was the most important predictor by a significant margin.

**Table 4.** A table depicting statistical parameter estimates from an elastic net logistic model (with clinical status as a binary dependent variable, and multiple brain age deltas [from different algorithms] and other covariates as independent variables). Standardized coefficients and standard errors are shown in different columns, with statistics for each independent variable shown in each row.

Elastic Net
| | $\beta$ (std. coef.) | Std. error |
| --- | --- | --- |
| **Real age** | 0.13370 | 0.00167 |
| **XGBoost delta** | 0.11000 | 0.00320 |
| **CAT12** | -0.03560 | 0.00180 |
| **DeepBrainNet delta** | 0.03300 | 0.00253 |
| **brainageR delta** | 0.01590 | 0.00203 |
| **Sex - male** | 0.00430 | 0.00137 |

## DISCUSSION

Through the analysis of multiple, large-scale, open-access neuroimaging datasets, we richly investigated critical elements of the calculation of brain age. Specifically, and with an eye toward applied research, we examined the reliability, noise sensitivity, and predictive power of three commonly used brain age algorithms (XGBoost, brainageR, and DeepBrainNet). Regarding reliability, we found all brain age algorithms were highly reliable. This was assessed through ICC and Bland-Altman metrics in samples of older and younger participants with repeated MRI scans. Related to brain age calculation and demographic variables of interest, there were strong correlations between these variables for each algorithm. Calculated brain age for each algorithm strongly tracked with chronological age across males and females (*r’s>*.*9*). Connected to image quality, all algorithms had significant negative correlations between brain age and image quality. Notably, this was with brain age, and not “brain age delta”. Correlations between brain age delta and image quality were modest for XGBoost, but near zero for brainageR and DeepBrainNet. Turning to clinical prediction, individual logistic models suggested that brain age delta was a significant predictor of dementia status. In models only including data derived from one brain age algorithm, XGBoost, brainageR, and DeepBrainNet all were individually related to MCI or Alzheimer’s disease status. In penalized regression models more suited to deal with collinear variables, brain age deltas derived from XGBoost were the strongest predictor of this clinical status.

Synthesizing across these different results, use of XGBoost may come with equal advantages and disadvantages. While all algorithms demonstrated excellent reliability as assessed by ICCs, it is notable that XGBoost had higher levels of bias (6.535%) in our younger sample of participants as assessed by Bland-Altman metrics. In the aggregate, this variation was small (-0.139 years) and that project (AOMIC) had a very narrow age range. However, in younger cohorts, this could lead to additional noise variance when examining this brain age algorithm in relation to important individual differences. Similarly, XGBoost’s brain age delta had a modest correlation with image quality. This will be important to consider when selecting brain age algorithms in different cohorts. If the population is likely to have high levels of movement (i.e., children; individuals with significant cognitive deficits and impulsivity), this could create additional noise, and cloud relations between brain age delta and other variables of interest. With all that duly noted, XGBoost was the most sensitive at differentiating individuals with significant clinical issues (i.e., MCI or Alzheimer’s disease status) across logistic and penalized models. Importantly, these models included a metric of image quality, suggesting that XGBoost’s brain age derivation explained important variance above and beyond this nuisance variable.

Thinking about our findings in relation to past reports, similar patterns have been noted individually for each algorithm regarding reliability and correlations with chronological age. Publications detailing the development of each algorithm have reported similar ICCs and correlations. Our project, however, is the first to examine these elements across multiple algorithms. While no algorithm that we investigated was superior on reliability metrics, it will be critical for future projects focused on novel metrics of brain age to compare performance to other algorithms as relative benchmarks. Put another way, publications that simply report metrics of their newly derived algorithm’s performance may be less useful to the field if this performance is not necessarily different or superior to other previously developed approaches. Interestingly, we did find better performance for XGBoost on differentiating MCI or Alzheimer’s disease status. Again, we believe this is the first publication to compare across algorithms to use the unique variance added by each algorithm. As noted in our results section, brainageR and DeepBrainNet were reasonably well correlated (r>.5) suggesting that these algorithms may be identifying similar patterns of advanced brain aging. It will again be critical for future projects focused on novel metrics of brain age to show that novel algorithms are identifying unique and additive variance in brain age.

Of note and important for future work is that we examined fairly significant clinical issues in thinking about prediction. There is ongoing work looking at different individual difference measures that span a more normative continuum of functioning. It could be particularly useful to see if brain ages from these different algorithms relate to these individual differences (e.g., stress exposure; general cognitive functioning; obesity, (Ronan et al., 2016; Shokri-Kojori et al., 2021). Similarly, it will be critical for future investigation to probe multimodal MRI calculations of brain age. All of the algorithms examined here focused on T1-weighted images, either processed in Freesurfer (XGBoost) or in original NIfTI format (brainageR; DeepBrainNet). More recent work has leveraged diffusion imaging, often in concert with T1-weighted images for prediction of brain age (Beck et al., 2021; Richard et al., 2018). It is likely that brain ages calculated through multimodal MRI and with multiple algorithms could be more powerful in explaining age-associated functional declines and disease.

While we believe we advanced applied understanding of brain age calculation, our work is not without limitations. First, all of our data is cross-sectional in nature, and it will be important to think about estimation and validation of different performance metrics in participants with repeated MRI scans separated by long periods of time. By seeing levels of within-and between-person change in relation to different algorithms, we may be able to derive a particularly powerful window into age-associated functional declines and disease, and different clinically relevant issues. Second, we did not specifically focus on variations in MRI scanners, instead pooling across scanning types. One cohort (HCP-Aging) had technically sophisticated MRI acquisition, potentially more sensitive than other “out of the box” neuroimaging scans. However, there were not notable patterns of differences in examining this cohort and the OASIS study that also contained similarly aged participants. Third, we tested three commonly used algorithms where code was publicly shared for mass implementation of brain age calculation. There are many in-press and preprinted manuscripts engineering new calculations of brain age. Such novel algorithms may exhibit superior performance and fewer limitations than the approaches we examined here. Fourth, we were unable to discern the factors that might be driving variations in algorithmic performance. Brain age calculated via XGBoost uses Freesurfer parcels in its brain age calculation, while brainageR and DeepBrainNet work off of less-processed NIfTI files. It may be possible to optimize elements of Freesurfer or other processes to improve different metrics of reliability and prediction. Connected to this, our team is particularly interested in the effects of image quality on brain age calculation and how to probe different datasets where repeated MRI scans are acquired from the same individuals but there is intentional variability in motion-related artifacts. Tackling these and other open questions related to brain age could significantly advance our understanding of healthy, as well as accelerated, aging processes.

Limitations notwithstanding, additional research on “*biological age*” is imperative. Richer information about the brain and brain aging could be important for those focused on age-related mortality and morbidity. Here, we provide important information about multiple brain age algorithms for researchers to consider when they deploy this emerging biomarker. Thoughtful consideration about reliability, noise tolerance, and predictive power will be critical when making decisions about different brain age algorithms, especially with an ever-growing landscape of potential ways to calculate this variable.

## Notes

https://www.oasis-brains.org/

https://nilab-uva.github.io/AOMIC.github.io/

https://www.humanconnectome.org/

### Competing Interest Statement

The authors have declared no competing interest.

