## Supplement for "Probing Multiple Algorithms to Calculate Brain Age: Examining Reliability, Relations with Demographics, and Predictive Power"

**Table S1,** *Caption: Summary of MRI data collection across the three datasets used in analysis. Columns contain information on age range, sample size, scanner, and additional details relevant to MRI scanning protocol.*

*MRI Data Collection*

| | **age range** | **sample size** | **scanner** | **additional details** |
|---|---|---|---|---|
| Amsterdam Open MRI Collection (AOMIC) | 19-26 | 928 | Phillips Intera 3T | TR/TE = 8.1ms/3.7ms<br>voxel size = 1 $mm^3$<br>matrix size = 64 x 64 |
| Open Access Series of Imaging Studies (OASIS) | 42-95 | 1098 | Siemens TIM Trio 3T | TR = 2.4 s<br>TE = 3.2 ms<br>vowel size = 1 $mm^3$<br>matrix size = 256 × 256 |
| Human Connectome Project in Aging (HCP-A) | 36-100 | 725 | Siemens Prisma 3T | TR = 2500/1000<br>TE = 1.8/3.6/5.4/7.2 ms<br>voxel size = 0.8 $mm^3$<br>matrix size = 320 × 300 × 208 |

**Table S2,** *Caption: Summary of correlations between variables calculating using the XGBoost model. The first column contains the two variables analyzed in that row, separated with a comma. The subsequent columns contain correlation coefficient r, t-statistic, and p-value.*

*XGBoost correlations*

| Variables | Correlation (r) | T-statistic | P-value |
|---|---|---|---|
| brain age, real age | 0.936 | 129.300 | 0.000 |
| brain age, real age (female) | 0.933 | 93.566 | 0.000 |
| brain age, real age (male) | 0.941 | 90.258 | 0.000 |
| brain age delta, real age | -0.776 | -59.615 | 0.000 |
| absolute brain age delta, real age | 0.543 | 31.355 | 0.000 |
| brain age delta, CAT12 score | 0.360 | 18.735 | 0.000 |
| absolute brain age delta, CAT12 score | -0.276 | -13.942 | 0.000 |
| real age, CAT12 Score | -0.381 | -19.964 | 0.000 |

**Table S3,** *Caption: Summary of correlations between variables calculating using the brainageR model. The first column contains the two variables analyzed in that row, separated with a comma. The subsequent columns contain correlation coefficient r, t-statistic, and p-value.*

*brainageR correlations*

| Variables | Correlation (r) | T-statistic | P-value |
|---|---|---|---|
| brain age, real age | 0.966 | 182.572 | 0.000 |
| brain age, real age (female) | 0.966 | 134.634 | 0.000 |
| brain age, real age (male) | 0.968 | 124.729 | 0.000 |
| brain age delta, real age | -0.225 | -11.216 | 0.000 |
| absolute brain age delta, real age | 0.307 | 15.651 | 0.000 |
| brain age delta, CAT12 score | -0.084 | -4.112 | 0.000 |
| absolute brain age delta, CAT12 score | -0.076 | -3.674 | 0.000 |
| real age, CAT12 Score | -0.458 | -24.995 | 0.000 |

**Table S4,** *Caption: Summary of correlations between variables calculating using the DeepBrainNet model. The first column contains the two variables analyzed in that row, separated with a comma. The subsequent columns contain correlation coefficient r, t-statistic, and p-value.*

*DeepBrainNet Correlations*

| Variables | Correlation (r) | T-statistic | P-value |
|---|---|---|---|
| brain age, real age | 0.960 | 166.890 | 0.000 |
| brain age, real age (female) | 0.958 | 121.240 | 0.000 |
| brain age, real age (male) | 0.963 | 116.068 | 0.000 |
| brain age delta, real age | -0.489 | -27.169 | 0.000 |
| absolute brain age delta, real age | 0.099 | 4.849 | 0.000 |
| brain age delta, CAT12 score | 0.033 | 1.622 | 0.105 |
| absolute brain age delta, CAT12 score | 0.033 | 1.626 | 0.104 |
| real age, CAT12 Score | -0.464 | -25.383 | 0.000 |

**Figure S1,** *Caption: Three scatterplots showing the relationship between real age (x-axis) and brain age delta (y-axis) on the AOMIC dataset. Each plot shows a different algorithm, in the order brainageR, DeepBrainNet, XGBoost. Female participants are indicated with red points and male participants are indicated with teal points.*

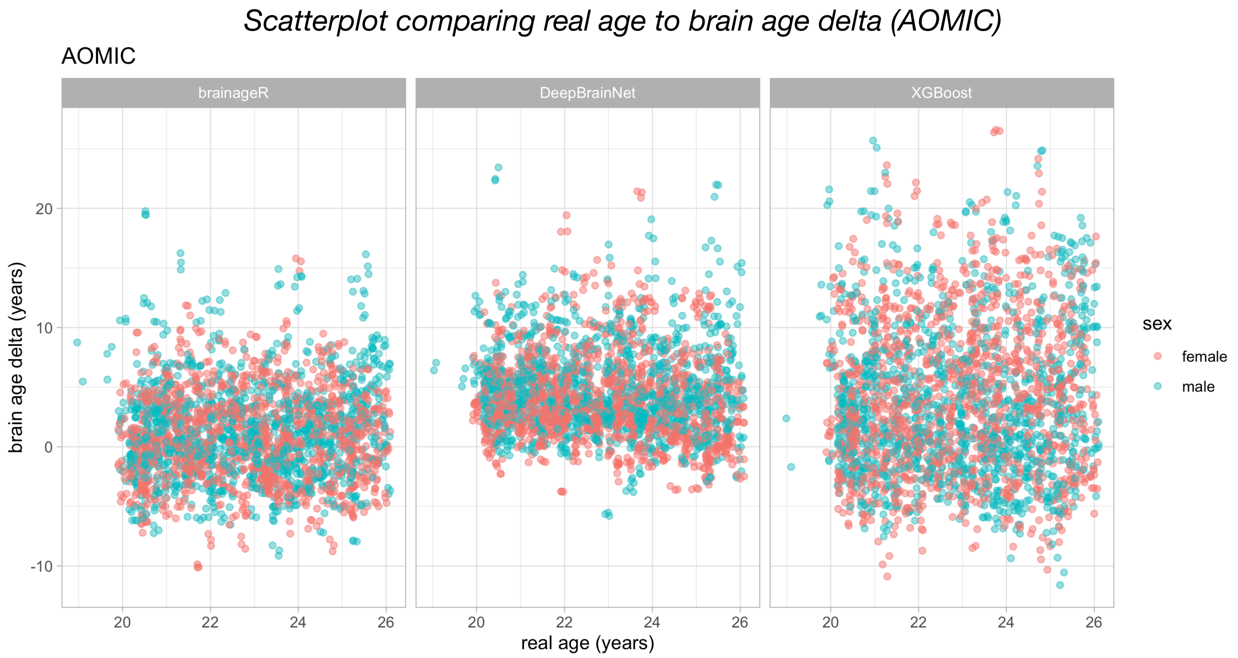


**Figure S2,** *Caption: Three scatterplots showing the relationship between real age (x-axis) and brain age delta (y-axis) on the HCP dataset. Each plot shows a different algorithm, in the order brainageR, DeepBrainNet, XGBoost. Female participants are indicated with red points and male participants are indicated with teal points.*

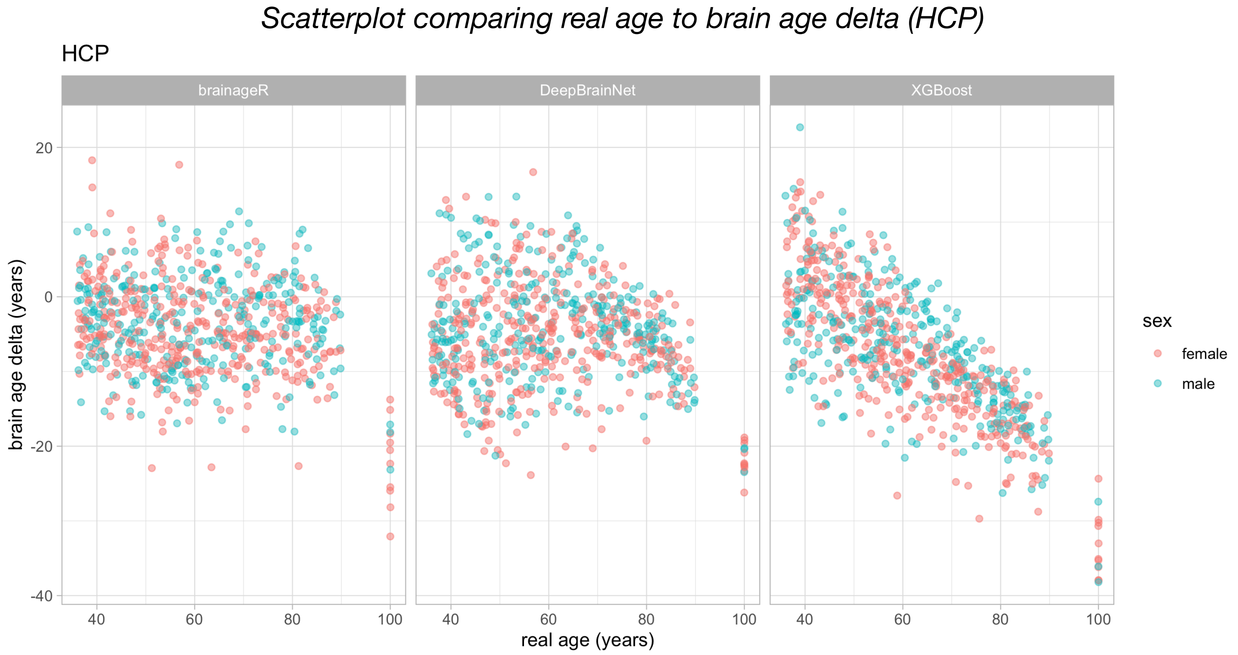

**Figure S3,** *Caption: Three scatterplots showing the relationship between real age (x-axis) and brain age delta (y-axis) on the OASIS dataset. Each plot shows a different algorithm, in the order brainageR, DeepBrainNet, XGBoost. Female participants are indicated with red points and male participants are indicated with teal points.*

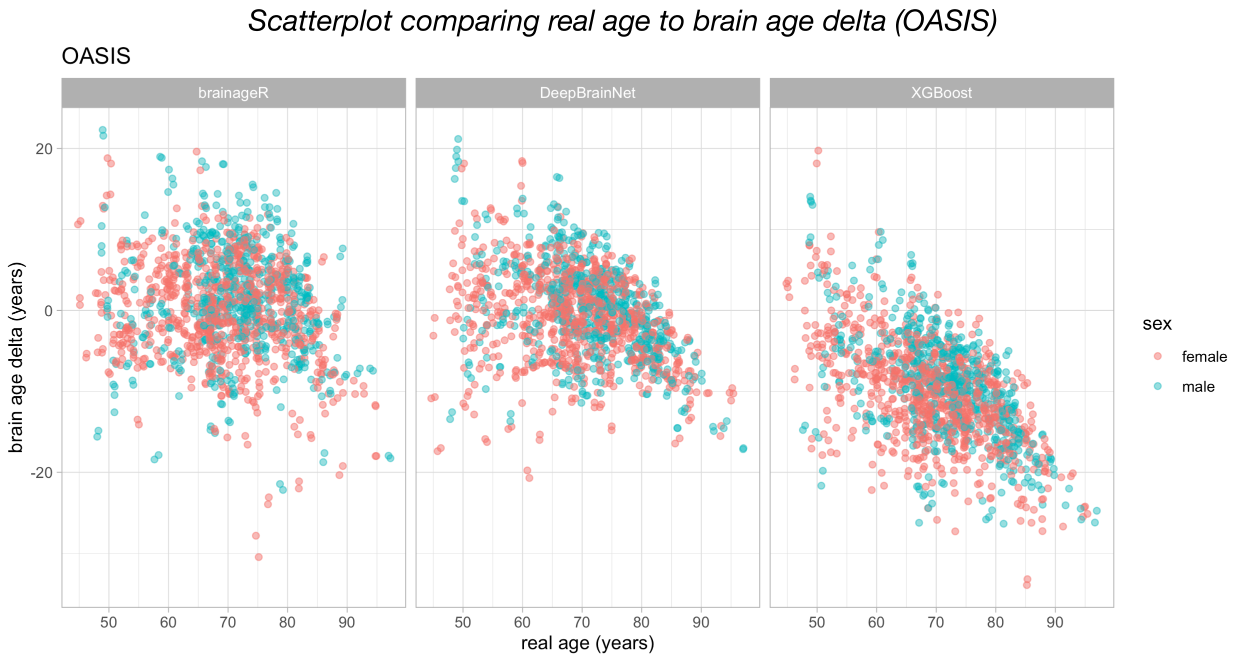
